# Analysis of a structured intronic region of the LMP2 pre-mRNA from EBV reveals associations with human regulatory proteins

**DOI:** 10.1101/405977

**Authors:** Nuwanthika Kumarasinghe, Walter N. Moss

## Abstract

Objective: The pre-mRNA of the Epstein–Barr virus LMP2 (<u>l</u>atent <u>m</u>embrane <u>p</u>rotein 2) has a region of unusual RNA structure that partially spans two consecutive exons and the entire intervening intron; suggesting RNA folding might affect splicing—particularly via interactions with human regulatory proteins. To better understand the roles of protein associations with this structured intronic region, we undertook a combined bioinformatics (motif searching) and experimental analysis (biotin pulldowns and RNA immunoprecipitations) of protein binding. Result: Characterization of the ribonucleoprotein composition of this region revealed several human proteins as interactors: regulatory proteins hnRNP A1 (<u>h</u>eterogeneous <u>n</u>uclear <u>r</u>ibo<u>n</u>ucleo<u>p</u>rotein <u>A1</u>), hnRNP U, HuR (<u>hu</u>man antigen <u>R</u>), and PSF (<u>p</u>olypyrimidine tract-binding protein-associated <u>s</u>plicing <u>f</u>actor), as well as, unexpectedly, the cytoskeletal protein actin. Treatment of EBV-infected cells with drugs that alter actin polymerization specifically showed marked effects on splicing in this region. This suggests a potentially novel role for nuclear actin in regulation of viral RNA splicing.

## INTRODUCTION

Epstein–Barr virus is a ubiquitous human herpes virus that infects ~95% of adults (1). EBV is implicated in various cancers (2-4) and autoimmune diseases (5). Host-virus interactions are crucial to infection and the emergence of disease (6). The precise mechanisms for EBV-implicated pathogenesis remain unclear, making molecular studies of this virus an active area of research. One particularly important area of study are the roles of RNA *inter*molecular (7-10) and *intra*molecular structures (11, 12), which have been found to be important to EBV infection. Intronic regions of the <u>l</u>atent <u>m</u>embrane <u>p</u>rotein 2 (LMP2) gene in <u>E</u>pstein–<u>B</u>arr <u>v</u>irus (EBV) were previously shown to be “hot spots” for stable and conserved RNA structure (12); suggesting potential roles in splicing regulation in LMP2, which is essential in establishing latent infection (13). In addition to the dominant isoforms of LMP2 (A and B), this gene possesses numerous, less abundant, minor isoforms generated from alternative splicing (14); a process known to be influenced by intronic RNA structure (6). The splice sites between LMP2 exons 7 and 8 are the only ones not utilized in alternative splicing and, interestingly, are the only splice sites to both be included within a single structural domain. Both are in helixes that form part of a 119 nt RNA structure that spans parts of both exons as well as the intervening intron (Fig. 1A). The unusual stability and conservation of this fold (12) suggests its importance to EBV, as do the presence of two unusual UU/UU internal “loop” motifs that are adjacent to each splice site (Fig. 1A). These motifs are rare, however, a single UU internal loop was previously found within the <u>i</u>ntron <u>s</u>plicing <u>s</u>ilencer (ISS) of HIV-1: an RNA structure important to splicing regulation (15). The sequence/structure of this ISS motif was found to associate with the human regulatory protein hnRNP A1 (<u>h</u>eterogeneous <u>n</u>uclear <u>r</u>ibo<u>n</u>ucleo<u>p</u>rotein <u>A1</u>). To understand how analogous interactions of human proteins with the LMP2 intronic structural region might be playing similar regulatory roles, we undertook a study to identify its <u>r</u>ibo<u>n</u>ucleo<u>p</u>rotein (RNP) composition.

**Figure 1.**
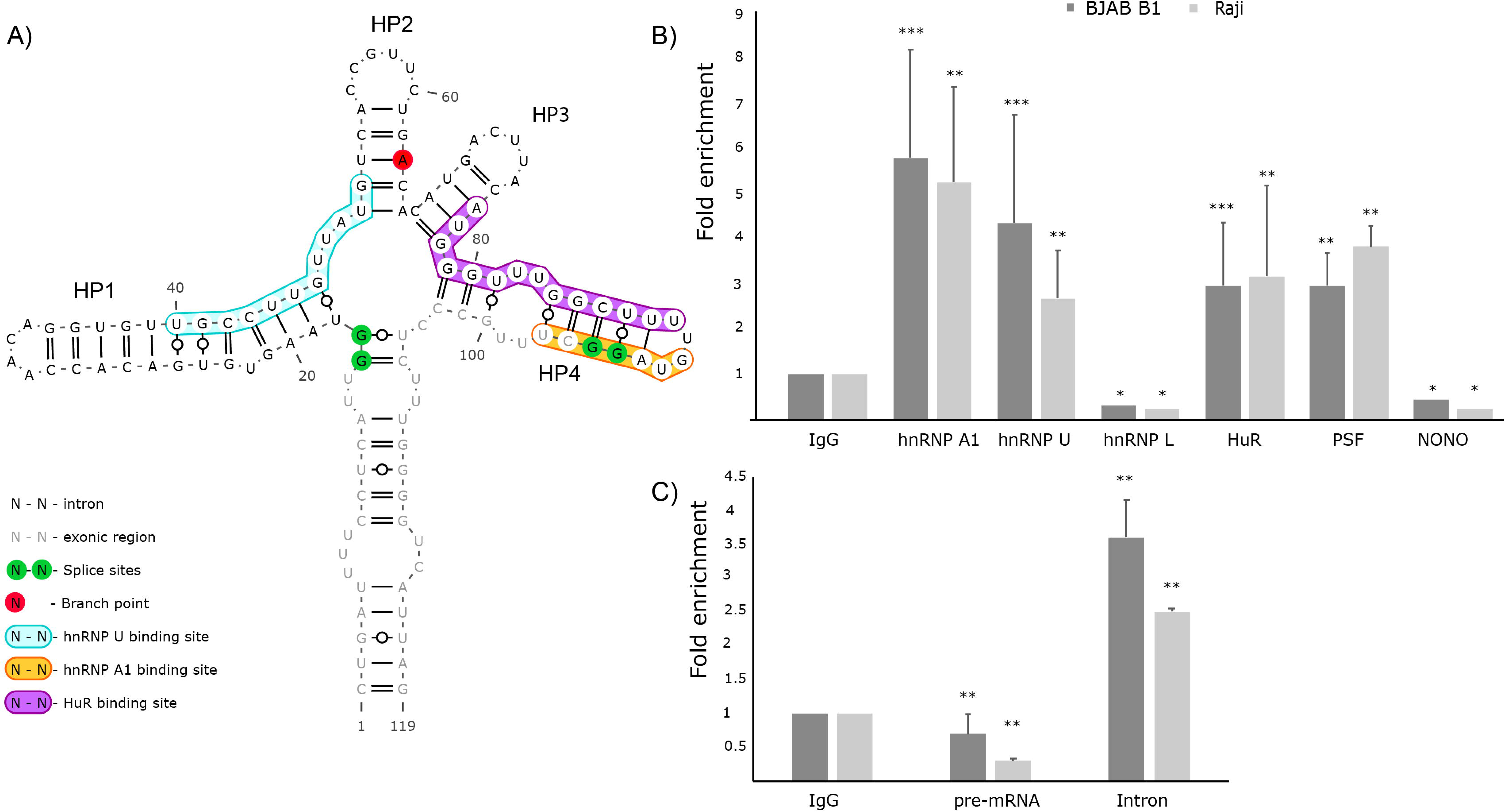
Validation of protein interactors. A) Secondary structure model of the intronic sequence. hnRNP A1, hnRNP U and HuR binding sites predicted by RBPmap are color coded. B) Fold enrichment of the LMP2 pre-mRNA following RIPs carried out with antibodies against hnRNP A1, hnRNP U, hnRNP L, HuR, PSF and NONO. C) Fold enrichment of pre-mRNA (junction spanning primers) and intron (internal primers) following RIPs with anti-actin antibody. Data represents the average (with standard deviation) of independent experiments all normalized to control RIP with IgG. *n=1; ** n=2; ***n=3 All primer sequences used for the experiments are included in Table S4.

## MAIN TEXT

To identify putative protein interactors we utilized the program RBPmap to scan the 119 nt LMP2 intronic RNA structured region for human regulatory protein binding motifs (16); complete results are in Table S1. Three predicted interacting proteins (Fig. 1A) were validated using <u>R</u>NA <u>i</u>mmuno<u>p</u>recipitations (RIPs; performed in two EBV-infected cell lines) followed by quantitative (q)PCR; the intronic region precipitated with HuR, hnRNP U and hnRNP A1 antibodies (Fig. 1B). HuR is involved in mRNA transport and affects post transcriptional modification via associations with additional regulatory proteins (17). hnRNP U, also termed <u>s</u>caffold <u>a</u>ttachment <u>f</u>actor <u>A</u> (SAF-A), interacts with a various pre-mRNAs, DNA elements with regions for nuclear matrix/scaffold attachment, and protein elements that include nuclear actin and the C-terminal domain (CTD) of RNA polymerase II. As mentioned previously, hnRNP A1 binds to a structured element within the HIV-1 ISS that, similar to EBV HP4, forms a hairpin that contains a UU internal loop motif. In HP4, however, the UU/UU motif is closer to the hnRNP A1 binding motif and the 3’ splice site, which in HP4 overlaps this motif (Fig. 1A). We also tested several other RNA-binding proteins that are predicted to be direct interactors or that might have protein-protein associations with HuR, hnRNP U or hnRNP A1. Although hnRNP L had a predicted binding site on the intronic structure, RIP data didn’t show any evidence of interaction (Fig. 1B). PSF (<u>p</u>olypyrimidine tract-binding protein-associated <u>s</u>plicing <u>f</u>actor) and p54 nrb/NONO (<u>N</u>on-POU domain-containing octamer-binding protein), which may interact with hnRNP U, were also tested. While NONO failed to show any significant enrichment compared to the IgG control, PSF (involved in mRNA processing (18)) precipitated the intronic region (Fig. 1B).

To identify additional protein interactors that may not directly associate with the intronic region, a pull-down assay was performed using biotinylated “bait” RNAs to precipitate interacting proteins from human B cell lysates. Various sized proteins were pulled down under different wash stringencies (Fig. S1); <u>M</u>ass <u>s</u>pectrometry (MS) was used to identify two bands that appeared in both medium and high stringency conditions. Consistent with *in silico* predictions and RIP, one band (around 37 kDa; Fig. S1) identified by MS was for hnRNP A1. Another band (37-50 kDa; Fig. S1) was found to be very prominent, even under high stringency washes. When identified via MS, the highest confidence result was for actin (MS results are in Table S2). RIP with anti-actin antibody confirmed that the intronic region was precipitated. Interestingly, only the intron between exon 7 and 8 could be amplified in the precipitated material (Fig. 1C). A 4-fold enrichment (vs. IgG control) was observed when qPCR was carried out with primers amplifying the intron; however, unlike the RIPs for other interactors that precipitated both pre-mRNA (Fig. 1B) and intron (Fig. S2), primers designed to amplify the intron-exon junction failed to show any significant enrichment in either BJAB-B1 or Raji cell lines. Actin should bind to both pre-mRNA and the intron sequence and the absence of pre-mRNA in actin pulldowns suggested that suggests that actin binding might promote splicing: e.g. the reaction occurs too quickly to capture the substrate with RIPs.

The association with actin was a surprise—nuclear actin is known to play roles in regulating transcription (19) and is hypothesized to affect mRNA maturation (20); however, no roles for actin in the transcription or splicing of viral RNAs were previously reported. To determine if nuclear actin could affect splicing of the LMP2 intronic region, we assessed the effects of dysregulated actin polymerization on viral splicing. A drug known to interfere with actin polymerization in live cells was tested. Latrunculin sequesters free monomeric “globular” G-actin, inhibiting actin polymerization (21). Consistent with a role for actin in stimulating splicing, the levels of unspliced transcripts remained relatively the same, while the spliced isoform levels showed a significant decrease over time via end-point RT-PCR (Fig. 2A) and RT-qPCR (Fig. 2B) in latrunculin treated BJAB-B1 cells.

**Figure 2.**
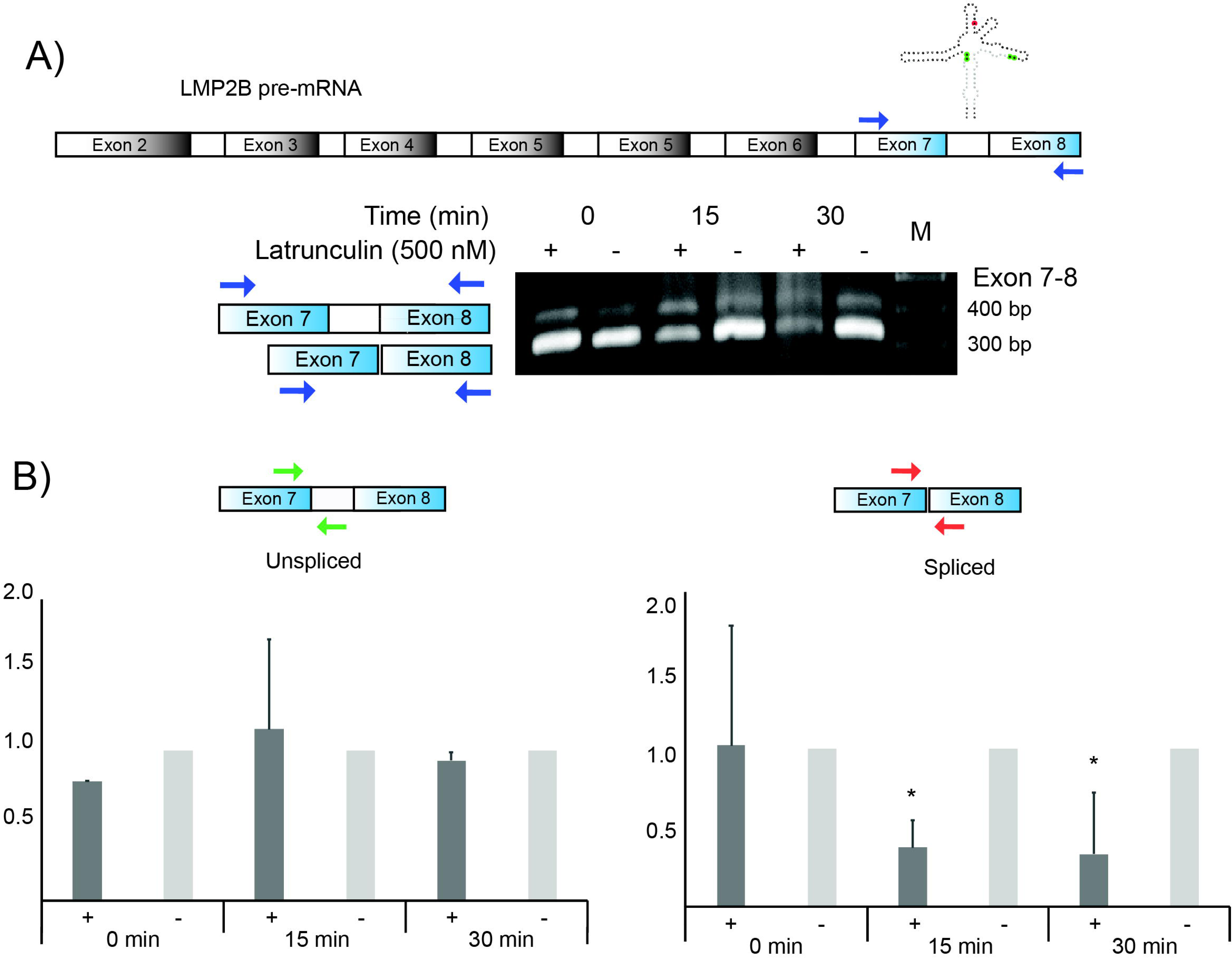
RT-PCR and qPCR analysis of spliced and unspliced transcripts following disruption of actin polymerization in BJAB B1 cells. A) A cartoon of the LMP2B with locations of exons RT-PCR primer sites and a model of the structured region is at the top. Below this are the results of RT-PCR analyses of spliced and unspliced transcripts in the presence of Latrunculin. B) Spliced and unspliced variants quantified by qPCR analysis. Data shown are first normalized to housekeeping gene HPRT and plotted as a fold difference compared to the control at each time point. All data represents the mean (with standard deviation) from two independent experiments. (*p < 0.05). All primer sequences used for the experiments are included in Table S4.

This preliminary observation points to a potential role for nuclear actin in the regulation of EBV mRNA processing. Interactions with regulatory proteins may also be playing roles here. For example, a STRING analysis of validated interactors (Table S3) finds an interaction network between hnRNP A1, hnRNP U and HuR, where HuR also has associations with actin. Additionally, hnRNP U was previously shown to interact with actin (18). The roles of these protein-protein associations, as well as the interactions with the LMP2 intron sequence/structure (e.g. the UU/UU motif on HP4) require additional study. Future work will elucidate these roles, determine how wide-spread are the roles played by actin in EBV mRNA processing (and beyond), and help resolve to what extent actin’s role is co-transcriptional vs. post-transcriptional.

## Materials and Methods

Detailed Materials and Methods can be found in Text S1.

## LIMITATIONS

The observations presented here are preliminary. Although we have validated several direct and indirect RNA-protein interactions in the intronic region, we are missing a map of the protein-protein interactions that form the RNP, we lack information on the exact roles of each interaction (in repression or stimulation of splicing) and the timing of these associations. We have suggestive results for actin, which point to a stimulatory roles for this association in splicing of this structured intronic region. However, we do not know how widespread are the effects of disrupting actin on splicing across EBV (and human) RNAs. These limitations, however, suggest many additional future analyses to parse out the roles of RNA structure and associations in LMP2 splicing and to better understand nuclear actin’s roles in regulating viral and host mRNA splicing.

**Abbreviations**
EBV: Epstein–Barr virus
LMP-2: latent membrane protein 2
ncRNA: non-coding RNA
qPCR: quantitative polymerase chain reaction
RBP: RNA binding protein
RNP: ribonucleoprotein
RT-PCR: reverse-transcriptase polymerase chain reaction

## Declarations

### Ethics approval and consent to participate

Not applicable

### Consent for publication

Not applicable

### Availability of data and material

Not applicable

## Authors’ contributions

NK: design and execution of experiments, data analysis/interpretation, and manuscript preparation. WNM: design and execution of data analysis/interpretation, manuscript preparation and final approval of the version to be published. All authors read and approved the final manuscript.

## Acknowledgements

We would like to thank all members of the Moss Lab, particularly Van Tompkins, for all the help with cell culture work and valuable suggestions. We would also like to thank Joel Nott at the Protein Facility of the ISU Office of Biotechnology for his help.

## Additional Files

**Text S1.** Detailed Materials and Methods.

**Figure S1.** Silver stained gels from biotin pulldowns. Bands P553-02 and P636-01B were identified as actin and hnRNPA with LC/MS-MS. M – Marker, L – lysate, FL – Full length 119 bp RNA construct of the structured intronic region, HP4 – truncated RNA construct containing only a 25 bp sub region HP4.

**Figure S2.** qPCR data showing the fold enrichment of the LMP2 intron following RIPs carried out with antibodies against hnRNP A1, hnRNP U, hnRNP L, HuR, PSF and NONO. Data represents the average (with standard deviation) of independent experiments all normalized to control RIP with IgG. *n=1; ** n=2; ***n=3

**Table S1.** RBPmap predictions of proteins predicted to bind motifs in the LMP2 structured intronic region.

**Table S2.** Mass spectrometry results on isolated silver stained bands (See Fig. S2) from biotin pulldown experiments.

**Table S3.** STRING analysis of validated protein interactors of the LMP2 structured intronic region.

**Table S4.** List of primers.

## References

1. Sitki-Green DL, Edwards RH, Covington MM, Raab-Traub N. 2004. Biology of Epstein-Barr virus during infectious mononucleosis. J Infect Dis 189:483–92.

2. Brady G, Macarthur GJ, Farrell PJ. 2008. Epstein-Barr virus and Burkitt lymphoma. Postgrad Med J 84:372–7.

3. Glaser SL, Lin RJ, Stewart SL, Ambinder RF, Jarrett RF, Brousset P, Pallesen G, Gulley ML, Khan G, O’Grady J, Hummel M, Preciado MV, Knecht H, Chan JK, Claviez A. 1997. Epstein-Barr virus-associated Hodgkin’s disease: epidemiologic characteristics in international data. Int J Cancer 70:375–82.

4. Gottschalk S, Rooney CM, Heslop HE. 2005. Post-transplant lymphoproliferative disorders. Annu Rev Med 56:29–44.

5. Toussirot E, Roudier J. 2008. Epstein-Barr virus in autoimmune diseases. Best Pract Res Clin Rheumatol 22:883–96.

6. Kuppers R. 2003. B cells under influence: transformation of B cells by Epstein-Barr virus. Nat Rev Immunol 3:801–12.

7. Lee N, Moss WN, Yario TA, Steitz JA. 2015. EBV noncoding RNA binds nascent RNA to drive host PAX5 to viral DNA. Cell 160:607–618.

8. Poling BC, Price AM, Luftig MA, Cullen BR. 2017. The Epstein-Barr virus miR-BHRF1 microRNAs regulate viral gene expression in cis. Virology 512:113–123.

9. Riley KJ, Rabinowitz GS, Yario TA, Luna JM, Darnell RB, Steitz JA. 2012. EBV and human microRNAs co-target oncogenic and apoptotic viral and human genes during latency. EMBO J 31:2207–21.

10. Tompkins VS, Valverde DP, Moss WN. 2018. Human regulatory proteins associate with non-coding RNAs from the EBV IR1 region. BMC Res Notes 11:139.

11. Cao S, Moss W, O’Grady T, Concha M, Strong MJ, Wang X, Yu Y, Baddoo M, Zhang K, Fewell C, Lin Z, Dong Y, Flemington EK. 2015. New Noncoding Lytic Transcripts Derived from the Epstein-Barr Virus Latency Origin of Replication, oriP, Are Hyperedited, Bind the Paraspeckle Protein, NONO/p54nrb, and Support Viral Lytic Transcription. J Virol 89:7120–32.

12. Moss WN, Steitz JA. 2013. Genome-wide analyses of Epstein-Barr virus reveal conserved RNA structures and a novel stable intronic sequence RNA. BMC Genomics 14:543.

13. Rovedo M, Longnecker R. 2007. Epstein-barr virus latent membrane protein 2B (LMP2B) modulates LMP2A activity. J Virol 81:84–94.

14. Concha M, Wang X, Cao S, Baddoo M, Fewell C, Lin Z, Hulme W, Hedges D, McBride J, Flemington EK. 2012. Identification of new viral genes and transcript isoforms during Epstein-Barr virus reactivation using RNA-Seq. J Virol 86:1458–67.

15. Jain N, Morgan CE, Rife BD, Salemi M, Tolbert BS. 2016. Solution Structure of the HIV-1 Intron Splicing Silencer and Its Interactions with the UP1 Domain of Heterogeneous Nuclear Ribonucleoprotein (hnRNP) A1. J Biol Chem 291:2331–44.

16. Paz I, Kosti I, Ares M, Jr., Cline M, Mandel-Gutfreund Y. 2014. RBPmap: a web server for mapping binding sites of RNA-binding proteins. Nucleic Acids Res 42:W361–7.

17. Papadodima O, Chatziioannou A, Patrinou-Georgoula M, Kolisis FN, Pletsa V, Guialis A. 2013. HuR-regulated mRNAs associated with nuclear hnRNP A1-RNP complexes. Int J Mol Sci 14:20256–81.

18. Kukalev A, Nord Y, Palmberg C, Bergman T, Percipalle P. 2005. Actin and hnRNP U cooperate for productive transcription by RNA polymerase II. Nat Struct Mol Biol 12:238–44.

19. Obrdlik A, Kukalev A, Louvet E, Farrants AK, Caputo L, Percipalle P. 2008. The histone acetyltransferase PCAF associates with actin and hnRNP U for RNA polymerase II transcription. Mol Cell Biol 28:6342–57.

20. Zheng B, Han M, Bernier M, Wen JK. 2009. Nuclear actin and actin-binding proteins in the regulation of transcription and gene expression. FEBS J 276:2669–85.

21. Peng GE, Wilson SR, Weiner OD. 2011. A pharmacological cocktail for arresting actin dynamics in living cells. Mol Biol Cell 22:3986–94.

